# Individual cortical neurons innervating large proportions of the neocortical areas in the MouseLight database

**DOI:** 10.1101/2025.10.22.683845

**Authors:** Szilvia Szeier, Emanuel Svensson, Ruben Wangel, Henrik Jörntell

**Affiliations:** Department of Experimental Medical Science, Lund University, Sweden

## Abstract

The general notion of the extent of cortical areal interconnectivity in the neuroscience community can in part be due to results from traditional neuroanatomical studies, which later insights have shown to systematically represent underestimates of the extent of the axonal arborizations. But any underestimate of this interconnectivity could in turn be a factor in strengthening the notion of functional localization in the cortex, i.e. the idea that there are circumscribed cortical areas with specific functions that do not to a large extent depend on information being processed in other such cortical areas. Recent advances in neuroanatomical techniques have greatly improved the possibilities to follow the axonal projections of individual neurons in full, and many reconstructed neurons are made available by the MouseLight database. We sampled individual cortical neurons with the majority of their axons within the neocortex and explored the extent of their axonal connections. We found that the efferent axon of individual cortical neurons can commonly cover as much as 30-40% of all types of cortical areas, with a coverage of up to 80% for a single neuron also being demonstrated. Combining the distributions of the axonal trees of more than one cortical neuron, we found that as few as three neurons within one area could reach 100% of the other cortical areas.

## 2 Introduction

Traditionally, neuroscience has strongly emphasized the importance of regional functional specialization in the neocortex. There has over the years indeed been results from several different methodologies which support that notion. However, recent studies which have not relied on ‘tip-of-the-iceberg’ effects but instead investigated the actual spread of information or activity across the cortex have suggested that the cortical network is much more globally integrated [1, 2, 3, 4, 5, 6, 7, 8, 9]. Even information about the specific quality of tactile or visual inputs is present in apparently any region of the neocortex [10, 11, 12, 13] and remote stroke-like lesions degrade the processing of tactile inputs in neurons in the primary somatosensory cortex (SOp) [14]. Cortical activation in a remote locations, as well as hippocampal output, also impact the neuronal interpretation/representation of tactile events in S1 neurons [15, 16]. All of these findings are at odds with the notion of the neocortex consisting of a set of functionally parcellated areas, beyond the fact the sensory information of different modalities enters the neocortex preferentially within limited specific cortical areas (and thereby the most easily detected effect when using ‘tip-of-the-iceberg’ methods).

Previous neuroanatomical studies, where the projections of axons from a particular cortical area for example, have to some extent rendered support for the functional localization case in the neocortex [17, 18]. However, axons are among the thinnest of the neural elements and the full extent of individual axons have always been difficult to trace with traditional neuroanatomical studies. More recently, methodological advances such as CLARITY [19] combined with laser scanning of fluorophore-labelled neurons within whole brain preparations appears to have improved on this shortcoming of the more traditional methods to trace individual axons [20]. The MouseLight project (https://www.janelia.org/project-team/mouselight) has accummulated a database of full anatomical reconstructions of individual neurons. Some neuroanatomical studies based on this database have indicated that the extent of connections made by individual corticocortical axons have previously been greatly underestimated [21], hence being in a position of providing a potential structural support for the observations that information in the neocortex is globally integrated. Studies of projection patterns from neurons in the prefrontal cortex are in support of this idea, where the combined projection pattern of in the order of 1000 prefrontal cortical neurons was shown to cover essentially the entire neocortex [22]. Here we wanted to explore the extent to which the above physiological observations could be explainable by pure structural information from the MouseLight project database. As the MouseLight project provides an open source database of many 100’s of cortical neurons, reporting the full extent of each axonal tree within the limits of the method, we utilized that database to explore how wide an extent of axonal branching across different cortical areas that each individual cortical neuron possessed.

Specifically, given that the database shows that individual cortical neurons can branch extensively within the cortex itself, we wanted to explore how many neurons located within one ‘area’ of the neocortex (i.e., isocortex) it would take to reach any other cortical area. We selected the four areas with the highest neuron density in the database and aimed to quantify the extent to which individual neurons’ axonal branches covered all other cortical areas, as well as the maximum coverage achievable by combining the axonal branches of multiple neurons. We find that individual cortical neurons can commonly cover as much as 30-40% of all cortical areas, with some individual neurons covering up to 80% of all cortical areas. Combining the distribution of the axonal tree of more than one cortical neuron located within the same cortical area, we found that as few as three neurons could reach 100% coverage, i.e. innervating all of the cortical areas.

## 3 Methods

All neuron data was downloaded from the MouseLight project at Janelia Research Campus (https://ml-neuronbrowser.janelia.org/, version 2022-02-28), which yielded 1227 neurons. Neurons are equipped with region identifiers for their soma, axon, and dendrite data points which are called AllenIDs. These AllenIDs correspond to mouse brain region labels within the Allen Mouse Brain Common Coordinate Framework (CCF) [23]. The CCF parcellated the mouse ‘isocortex’ (neocortex) into a total of 43 areas. We defined isocortex from the complete ontology of CCF (found in the supplementary information in [23]) as those structures which were within isocortex (their “structure id path” started with the isocortex “structure id path) and were a summary structure (“‘Summary Structure’ Level for Analysis” contained “Y”). The MouseLight project region labels further divide isocortical areas into more specific categories, e.g. “Primary motor area” into “Primary motor area, Layer 1”, “Primary motor area, Layer 2/3”, etc. As we were interested in the 43 isocortical regions defined by [23], any subdivisions of these regions were assumed to be part of that region. A full list of isocortical area abbreviations can be found in Table 1 and their specifications (name, abbreviation, path in tree structure, AllenIDs merged into that group) are provided in Supplementary Table 1. We applied this definition of isocortex to filter the entire set of neurons, selecting those with their soma located in one of these regions, resulting in a total of 450 isocortical neurons.

**Table 1:**
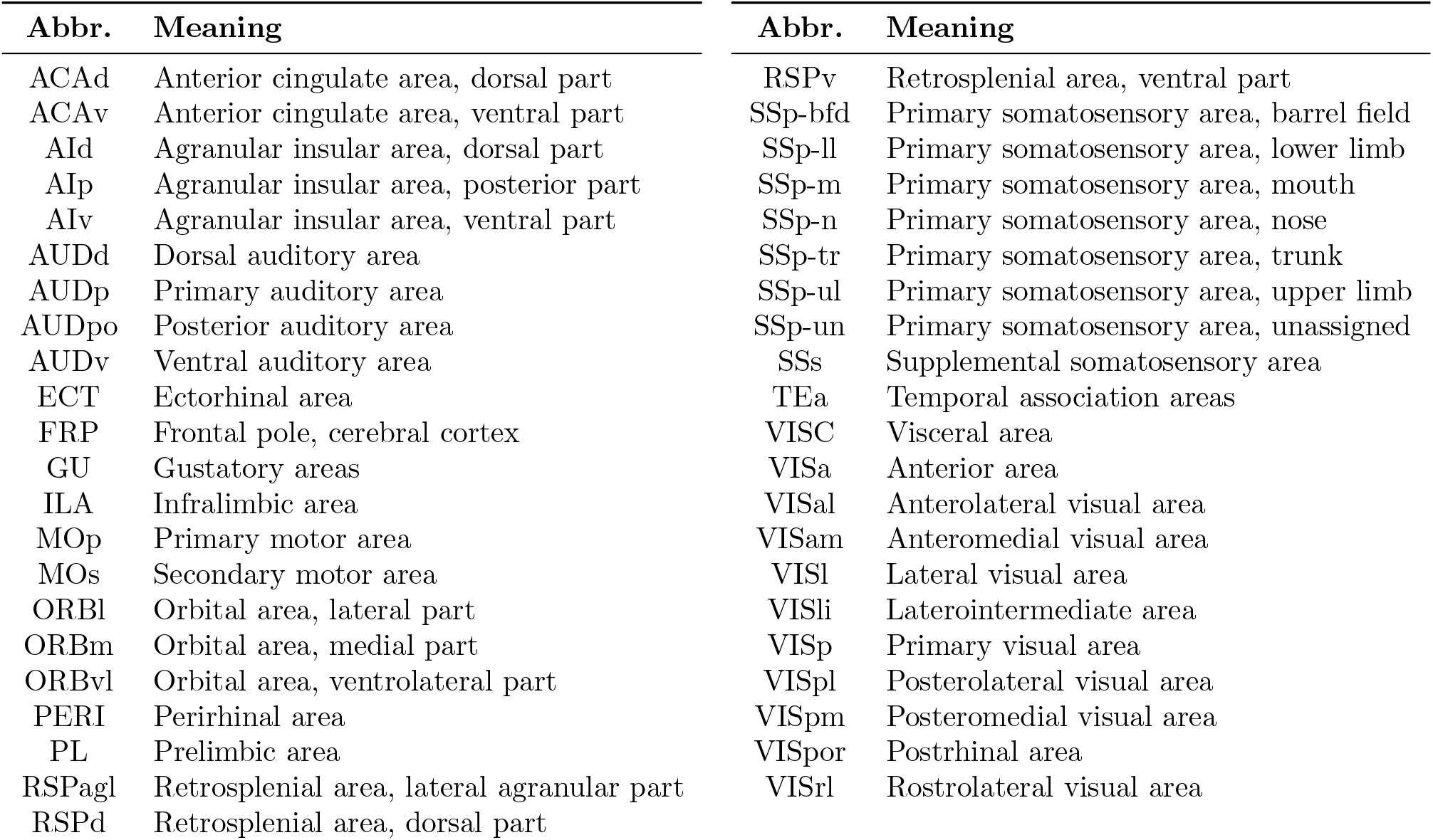
List of isocortical regions defined by CCF [23] and their abbreviations (abbr.).

For further investigations, we used the isocortical areas which contained the highest numbers of neurons. These regions were anterior cingulate area, dorsal part (ACAd), primary motor area (MOp), secondary motor area (MOs), and primary visual area (VISp) with 30, 48, 296, and 16 neurons respectively. We were interested in those neurons which predominantly maintain their axonal projections within the isocortex. Since each neuron is associated with AllenIDs (area labels) for all of its axon data points, we could calculate the proportion of a neuron’s axon located within the isocortex by dividing the number of AllenIDs located in the 43 isocortical regions by the total axon length. Neurons which had 60% or more of their axonal projections within the isocortex were further examined in our study. A full list of the neurons used can be found in Supplementary Table 2. For these neurons, we then determined the proportion of isocortical areas they covered. An area was considered covered if the neuron had at least one axonal data point within it. The proportion was calculated as the number of isocortical regions covered divided by the total number of regions (43).

To quantify the extent to which a population of neurons within a specific area could innervate other isocortical areas, we identified the minimum number of neurons needed to cover as many areas as possible (at most 43). Starting with the neuron that covers the most isocortical areas, we iteratively added neurons that maximally increased the total coverage of neocortical areas at each step. This process continued until no further increases in the coverage occurred.

## 4 Results

We used the neuron reconstructions from the MouseLight project. The database contains 1227 neurons out of which 450 had their somata located in one of the isocortical areas defined by Allen Mouse Brain Common Coordinate Framework (CCF) [23] (Figure 1A). The example pyramidal neuron (Figure 1B) demonstrates that individual neurons could have a wide span in the telencephalon. It’s worth noting that the axon branched extensively within the ipsilateral hemisphere and also spread significantly within the contralateral hemisphere. Although not all of these branches were within the cortex, most were. The database provided the location of each axon data point, allowing us to count the number of cortical areas innervated by the axon of each individual neuron.

**Figure 1:**
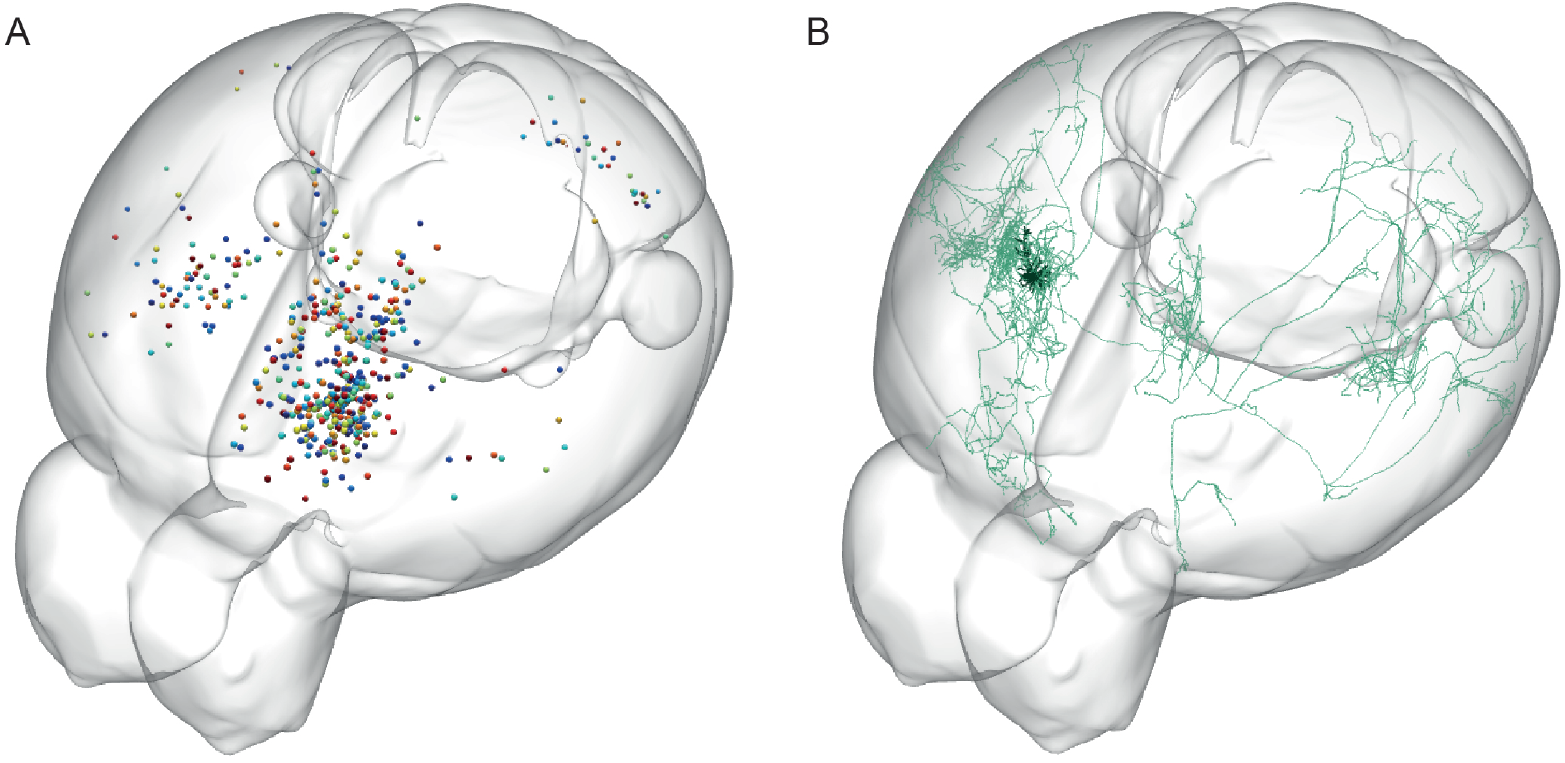
Visualization of all neurons with somata in the “Isocortex” in the MouseLight Browser (A), and an example axonal arborization of a single neuron AA0100 (B) shown in green, the neuron and its dendrites are in black. Images taken from the MouseLight database https://ml-neuronbrowser.janelia.org/.

The regional distribution of all isocortical somata included in this study can be seen in Figure 2. There were 43 isocortical areas defined in the CCF, but not all of them contained somata as retrieved by our search terms from the MouseLight project. Most neurons were located in MOs, secondary motor area, whereas also the primary motor area (MOp), the anterior cingulate area, dorsal part (ACAd), and the primary visual area (VISp) contained more than 15 neurons. We focused our investigation to these neurons.

**Figure 2:**
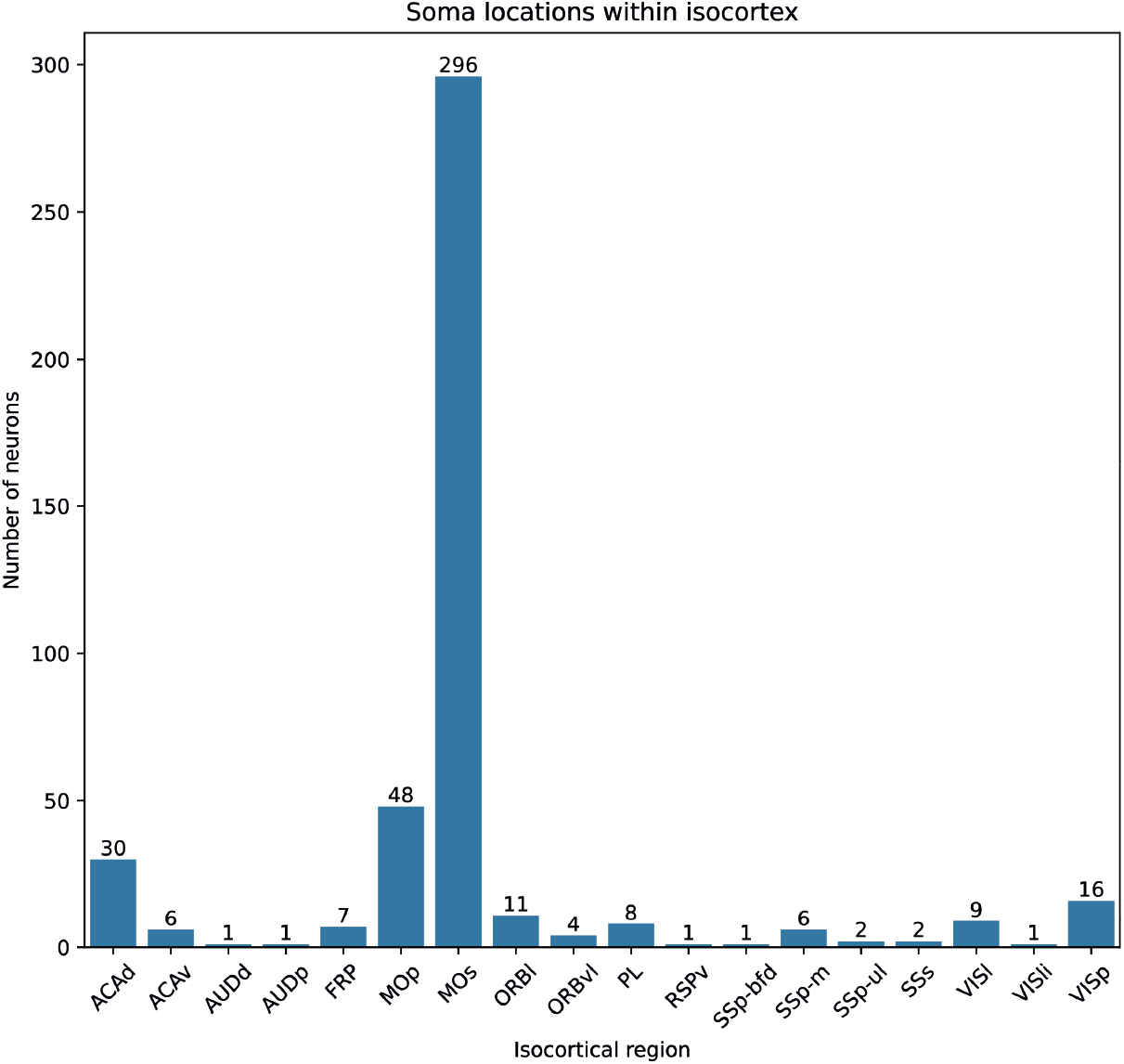
Distribution of the soma Allen IDs in the data set when queried for “isocortex” in the Neuron Browser at https://ml-neuronbrowser.janelia.org/. We selected the primary and secondary motor areas (MOp and MOs) along with anterior cingulate area, dorsal part (ACAd) and primary visual area (VISp), which contained the highest number of neurons. A full list of the abbreviations is found in Table 1.

For the neurons within each of the selected areas, we calculated the proportions of the axon data points that remained within the neocortex (Figure 3). In each area there were neurons that sent their axons exclusively outside the neocortex, and there were neurons that sent their axons exclusively within the neocortex, and then all kinds of mixtures in between. We limited our investigation to neurons with at least 60% of their axon data points within the neocortex.

**Figure 3:**
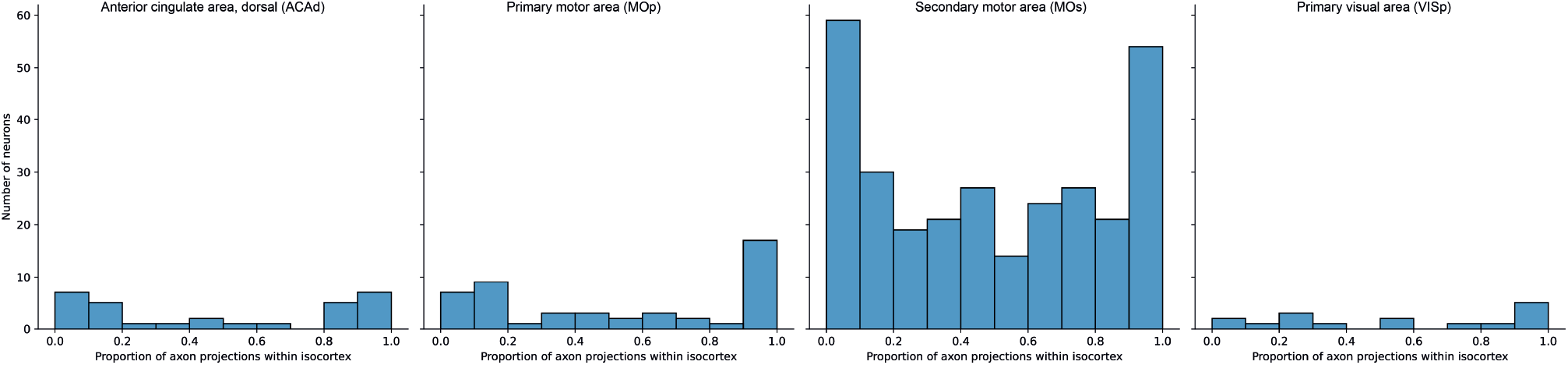
The selection of the neurons in the isocortical areas with 60% or more of their axon data points within the isocortex. This resulted in 13, 23, 126, 7 neurons for ACAd, MOp, MOs, VISp respectively.

The number of neurons which have at least 60% of their axon data point within the neocortex were 13, 23, 126, 7 neurons for ACAd, MOp, MOs, VISp respectively. Figure 4 illustrates the proportion of neocortical areas covered by each of these individual neurons. Quite remarkably, there were many neurons whose axon covered around 50% or even more of the 43 cortical areas.

**Figure 4:**
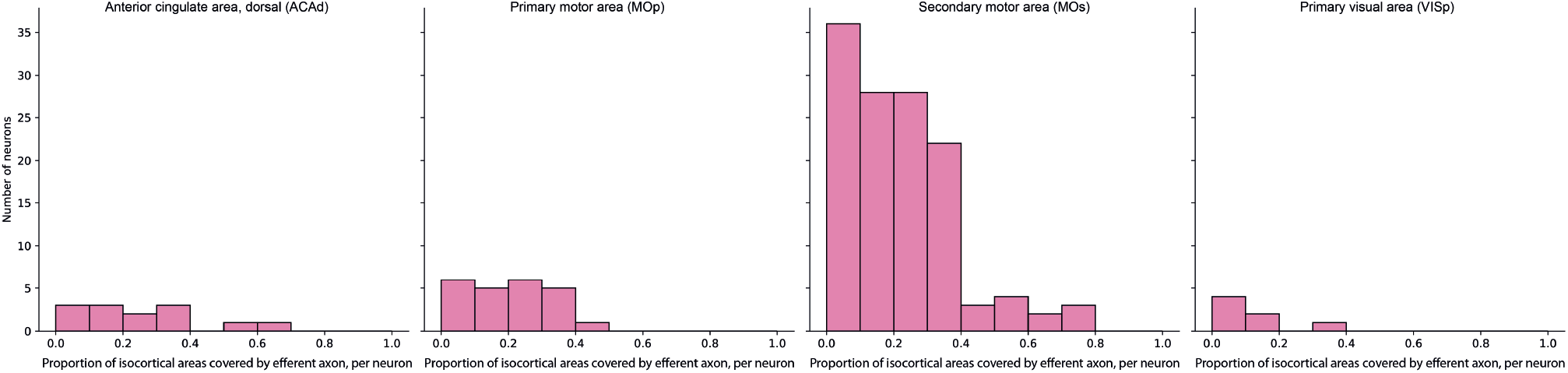
The proportion of isocortical areas covered by individual neurons which were defined to predominantly project within the neocortex in the highest neuron density isocortical regions (ACAd, MOp, MOs, VISp).

We also wanted to know, for each cortical area, if we picked a number of different neurons, how much more of neocortical areas could then be covered? This would basically answer the question if neurons within the same area tended to have similar or dissimilar projection patterns in their axonal branching patterns to other cortical areas. For each area, we picked the ‘best’ first neuron, and then we picked the second neuron as the one that would increase the proportion of neocortical areas covered the most, and so on for the third neuron, etc. Figure 5 shows that a third neuron still added substantial coverage, but also that as few as 3 neurons were enough to reach very high coverage of all the cortical areas. Then the coverage appeared to level out, at least for the limited number of neurons that the database contained. For ACAd we could reach 80% coverage of all neocortical areas with 3 neurons; MOp 67% with 4; MOs 100% with 3; VISp 37% with 2.

**Figure 5:**
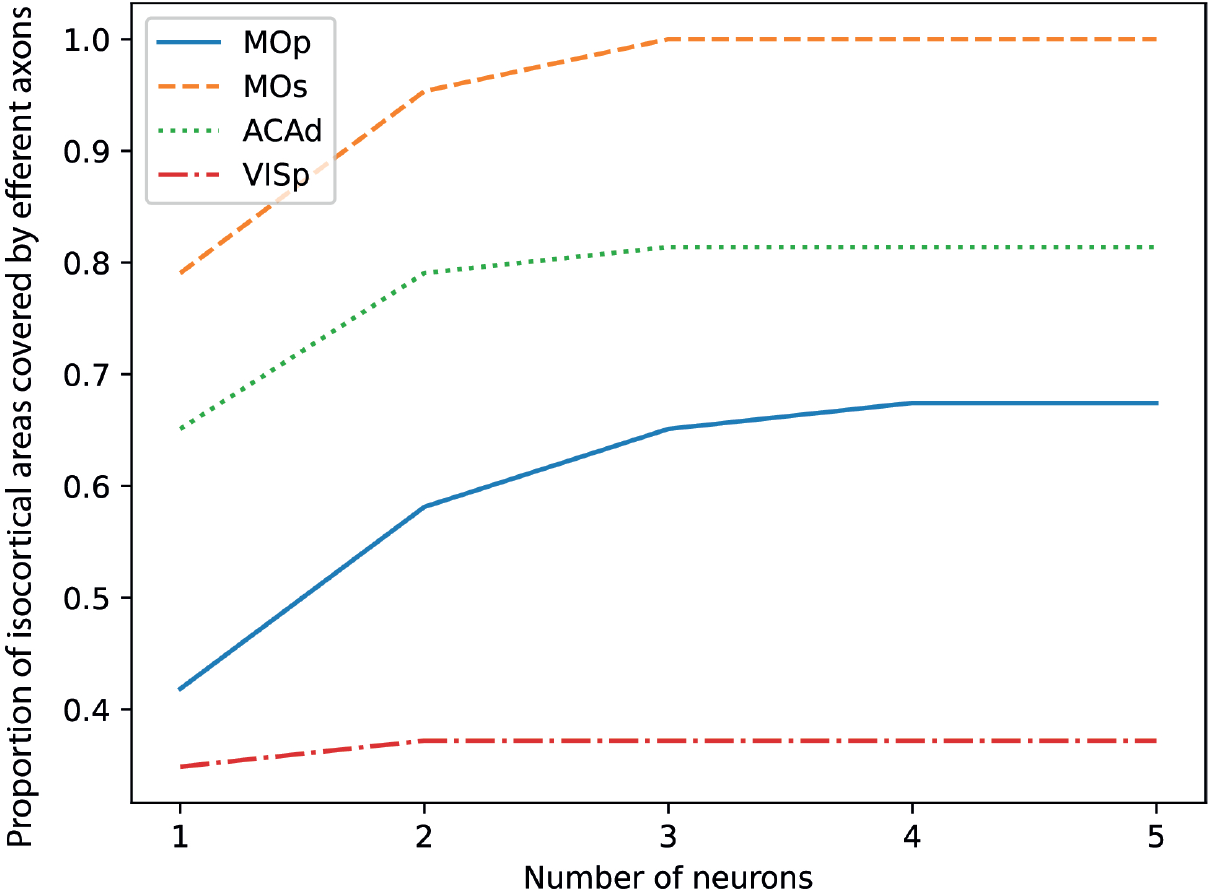
The minimal number of neurons needed to reach the maximal possible coverage. The individual neurons are [AA0034, AA0658, AA0002, AA0003] for MOp, [AA0100, AA0096, AA0332] for MOs, [AA0243, AA0098, AA0775] for ACAd, and [AA0638, AA0027] for VISp.

The axons of the neurons in VISp extended across AUDd, AUDp, AUDv, RSPagl, RSPd, SSp-bfd, SSs, TEa, VISa, VISal, VISam, VISl, VISli, VISp, VISpm, and VISrl. The Allen CCF classifies almost 25% of the cortical areas as being visual related (Table 1), which is surprising for an animal that is not supposed to rely so much on vision. It would seem likely that perhaps many of these areas that are carrying the ‘VIS*’ label are not directly or strictly related to visual processing. However, for the cortical areas covered by axons from the neurons located in VISp, 50% were ‘VIS*’ related, whereas among the remaining 50%, VISp neurons sent their axons to as remote locations as different subdivisions of the auditory cortex, as well as to secondary somatosensory cortex, retrosplenial areas and temporal association areas.

This result (Figure 5) is important because it suggests, in principle, that nearly every cortical area could potentially be covered by just a few neurons from any other individual cortical area. The primary visual cortex was the least extensively interconnected area. But then it should be recalled that there were only 16 neurons in total that were located in this area in the database. It is likely that if a more extensive sampling of neurons from this area had been available, it would contain neurons with more richly divergent connections, similar to those of neurons located in MOs, ACAd, and MOp.

## 5 Discussion

Our findings, based on the MouseLight database, suggest that neocortical neurons are extremely interconnected to the extent which could potentially substantially weaken the concept of functional localization within the mouse neocortex. Even though the dataset included only 450 isocortical neurons, far fewer than the actual count of more than 5 million cortical neurons in the mouse [24], we still found that, within a single functional region of at least 100,000 neurons (using the Allen definition of the cortex as consisting of 43 isocortical areas), just 3 neurons could be sufficient to reach all other neocortical areas.

It should be noted that because of this extreme subsampling of neurons in the MouseLight database, the coverage of neocortical areas can only represent an underestimate of the true extent of the axonal trees, as it is unlikely that the database represents the most extreme cases of corticocortical divergence of single neurons. Furthermore, the data collection procedure relied on fluorescent signals. Although a signal amplification method was employed [21], it’s possible that the fluorescent reporter (i.e. the fluorophore) did not always reach every part of each individual neuron.

We based our coverage estimates on the assumption that if a neuron had an axon data point in a given region then it also forms at least one synapse there. This assumption was motivated by that Winnubst et al. state that “although our pipeline does not explicitly detect synapses, axonal arborizations contain an approximately constant density of synapses, and axonal length is therefore a convenient surrogate for interareal connectivity” [21].

Finally, the data reported here of course only represent the monosynaptic connectivity. Much more extensive, and dense, interconnectivity can be expected if one also include disynaptic connectivity, not the least via the cortico-thalamo-cortical connections (see discussion in [11]). But whereas more indirect connections could always be speculated to be under different forms of conditional gating control, the very wide extent of monosynaptic connections between cortical areas (also shown for neurons in the prefrontal cortex [22]) makes it unavoidable that there is an extensive, non-suppressible information exchange between all of what has traditionally been referred to as functionally specialized regions of the cortex.

## References

[1] Isabelle Ferezou, Florent Haiss, Luc J Gentet, Rachel Aronoff, Bruno Weber, and Carl CH Petersen. Spatiotemporal dynamics of cortical sensorimotor integration in behaving mice. Neuron, 56(5): 907–923, 2007.

[2] Ron D Frostig, Ying Xiong, Cynthia H Chen-Bee, Eugen Kvašňák, and Jimmy Stehberg. Large-scale organization of rat sensorimotor cortex based on a motif of large activation spreads. Journal of Neuroscience, 28(49): 13274–13284, 2008.

[3] Kai-Ming G Fu, Taylor A Johnston, Ankoor S Shah, Lori Arnold, John Smiley, Troy A Hackett, Preston E Garraghty, and Charles E Schroeder. Auditory cortical neurons respond to somatosensory stimulation. Journal of Neuroscience, 23(20): 7510–7515, 2003.

[4] Asif A Ghazanfar and Charles E Schroeder. Is neocortex essentially multisensory? Trends in cognitive sciences, 10(6): 278–285, 2006.

[5] Sayaka Hihara, Miki Taoka, Michio Tanaka, and Atsushi Iriki. Visual responsiveness of neurons in the secondary somatosensory area and its surrounding parietal operculum regions in awake macaque monkeys. Cerebral Cortex, 25(11): 4535–4550, 2015.

[6] Georg B Keller, Tobias Bonhoeffer, and Mark Hübener. Sensorimotor mismatch signals in primary visual cortex of the behaving mouse. Neuron, 74(5): 809–815, 2012.

[7] Umberto Olcese, Giuliano Iurilli, and Paolo Medini. Cellular and synaptic architecture of multisensory integration in the mouse neocortex. Neuron, 79(3): 579–593, 2013.

[8] Ede A Rancz, Javier Moya, Florian Drawitsch, Alan M Brichta, Santiago Canals, and Troy W Margrie. Widespread vestibular activation of the rodent cortex. Journal of Neuroscience, 35(15): 5926–5934, 2015.

[9] Aman B Saleem, Aslı Ayaz, Kathryn J Jeffery, Kenneth D Harris, and Matteo Carandini. Integration of visual motion and locomotion in mouse visual cortex. Nature neuroscience, 16(12): 1864–1869, 2013.

[10] Jonas MD Enander and Henrik Jörntell. Somatosensory cortical neurons decode tactile input patterns and location from both dominant and non-dominant digits. Cell reports, 26(13): 3551–3560, 2019.

[11] Jonas MD Enander, Anton Spanne, Alberto Mazzoni, Fredrik Bengtsson, Calogero Maria Oddo, and Henrik Jörntell. Ubiquitous neocortical decoding of tactile input patterns. Frontiers in cellular neuroscience, 13:140, 2019.

[12] Clara Genna, Calogero M Oddo, Alberto Mazzoni, Anders Wahlbom, Silvestro Micera, and Henrik Jörntell. Bilateral tactile input patterns decoded at comparable levels but different time scales in neocortical neurons. Journal of Neuroscience, 38(15): 3669–3679, 2018.

[13] Sofie S Kristensen, Kaan Kesgin, and Henrik Jörntell. High-dimensional cortical signals reveal rich bimodal and working memory-like representations among s1 neuron populations. Communications biology, in press, 2024.

[14] Anders Wahlbom, Jonas MD Enander, Fredrik Bengtsson, and Henrik Jörntell. Focal neocortical lesions impair distant neuronal information processing. The Journal of physiology, 597(16): 4357–4371, 2019.

[15] Leila Etemadi, Jonas MD Enander, and Henrik Jörntell. Remote cortical perturbation dynamically changes the network solutions to given tactile inputs in neocortical neurons. Iscience, 25(1), 2022.

[16] Leila Etemadi, Jonas MD Enander, and Henrik Jörntell. Hippocampal output profoundly impacts the interpretation of tactile input patterns in si cortical neurons. iScience, 26(6), 2023.

[17] JW Neal, RCA Pearson, and TPS Powell. The ipsilateral corticocortical connections of area 7 with the frontal lobe in the monkey. Brain research, 509(1): 31–40, 1990.

[18] James W Lewis and David C Van Essen. Corticocortical connections of visual, sensorimotor, and multimodal processing areas in the parietal lobe of the macaque monkey. Journal of Comparative Neurology, 428(1): 112–137, 2000.

[19] Kwanghun Chung, Jenelle Wallace, Sung-Yon Kim, Sandhiya Kalyanasundaram, Aaron S Andalman, Thomas J Davidson, Julie J Mirzabekov, Kelly A Zalocusky, Joanna Mattis, Aleksandra K Denisin, et al. Structural and molecular interrogation of intact biological systems. Nature, 497(7449): 332–337, 2013.

[20] Charles R Gerfen, Michael N Economo, and Jayaram Chandrashekar. Long distance projections of cortical pyramidal neurons. Journal of neuroscience research, 96(9): 1467–1475, 2018.

[21] Johan Winnubst, Erhan Bas, Tiago A Ferreira, Zhuhao Wu, Michael N Economo, Patrick Edson, Ben J Arthur, Christopher Bruns, Konrad Rokicki, David Schauder, et al. Reconstruction of 1,000 projection neurons reveals new cell types and organization of long-range connectivity in the mouse brain. Cell, 179(1): 268–281, 2019.

[22] Le Gao, Sang Liu, Lingfeng Gou, Yachuang Hu, Yanhe Liu, Li Deng, Danyi Ma, Haifang Wang, Qiaoqiao Yang, Zhaoqin Chen, et al. Single-neuron projectome of mouse prefrontal cortex. Nature Neuroscience, 25(4): 515–529, 2022.

[23] Quanxin Wang, Song-Lin Ding, Yang Li, Josh Royall, David Feng, Phil Lesnar, Nile Graddis, Maitham Naeemi, Benjamin Facer, Anh Ho, et al. The allen mouse brain common coordinate framework: a 3d reference atlas. Cell, 181(4): 936–953, 2020.

[24] Suzana Herculano-Houzel, Bruno Mota, and Roberto Lent. Cellular scaling rules for rodent brains. Proceedings of the National Academy of Sciences, 103(32): 12138–12143, 2006.

